# Shared and unique responses in the microbiome of allopatric lizards reared in a standardized environment

**DOI:** 10.1101/2022.09.26.509558

**Authors:** Braulio A. Assis, Terrence H. Bell, Heather I. Engler, William L. King

## Abstract

The gut microbiome can influence host fitness and, consequently, the ecology and evolution of natural populations. Microbiome composition can be driven by environmental exposure but also by the host’s genetic background and phenotype. To contrast environmental and genetic effects on the microbiome we leverage preserved specimens of eastern fence lizards from allopatric lineages east and west of the Mississippi River but reared in standardized conditions. Bacterial composition was indistinguishable between lineages but responded significantly to host age – a proxy for environmental exposure. This was accompanied by a continuous decrease in bacterial diversity in both lineages, partially driven by decreasing evenness seen only in western lizards. These findings indicate that longer exposure to a homogeneous habitat may have a depreciating effect on microbiome diversity in eastern fence lizards, a response shared by both lineages. We highlight the importance of such effects when extrapolating patterns from laboratory experiments to the natural world.

## Introduction

Animals establish vital relationships with the microbial colonizers of the gastrointestinal tract (*i*.*e*. the gut microbiome) (Ley et al. 2008). Gut microbial composition is linked to multiple fitness-related traits such as nutrient assimilation (Kreznar et al. 2017; Newsome et al. 2020; Parata, Mazumder, et al. 2020), immune function (Round and Mazmanian 2009; Clavel et al. 2017), and behavior (Sylvia and Demas 2018), having substantial impacts on the ecology and evolution of natural populations (Macke et al. 2017; Gould et al. 2018). However, gut microbiomes can be shaped by factors that range from the host’s ecology (*e*.*g*., habitat, diet, phenology) (Wu et al. 2018; Youngblut et al. 2019; Parata, Nielsen, et al. 2020) to host genetics (Bonder et al. 2016), possibly with interactive effects. While controlled laboratory experiments eliminate environmental variation and isolate variables of interest, the effects of long-term standardized environments on host microbiomes require further investigation. This can be compounded if sampling involves hosts of distinct genetic backgrounds that show unique microbiome responses to the environment, compromising the detectability of biological relationships.

Here, we characterized the cloacal bacterial microbiome of spiny lizards (*Sceloporus spp,* figure 1.) from two distinct lineages that were reared in standardized conditions. To that end, we leverage preserved specimens that were part of a parallel experiment. Eggs were laid and hatched in an animal housing facility by wild females obtained from populations East and West of the Mississippi River. Phylogenetic analyses suggest restricted gene flow between the two populations (Leaché 2009; Wiens et al. 2010), despite being ecologically equivalent. Our objective was to assess whether population history will underlie either shared or unique microbiome responses in the face of identical environmental conditions throughout a host’s ontogeny (Spor et al. 2011). With that, we hope to gain better understanding of the roles that genetic background and environmental effects have on gut microbial compositions.

**Figure 1:**
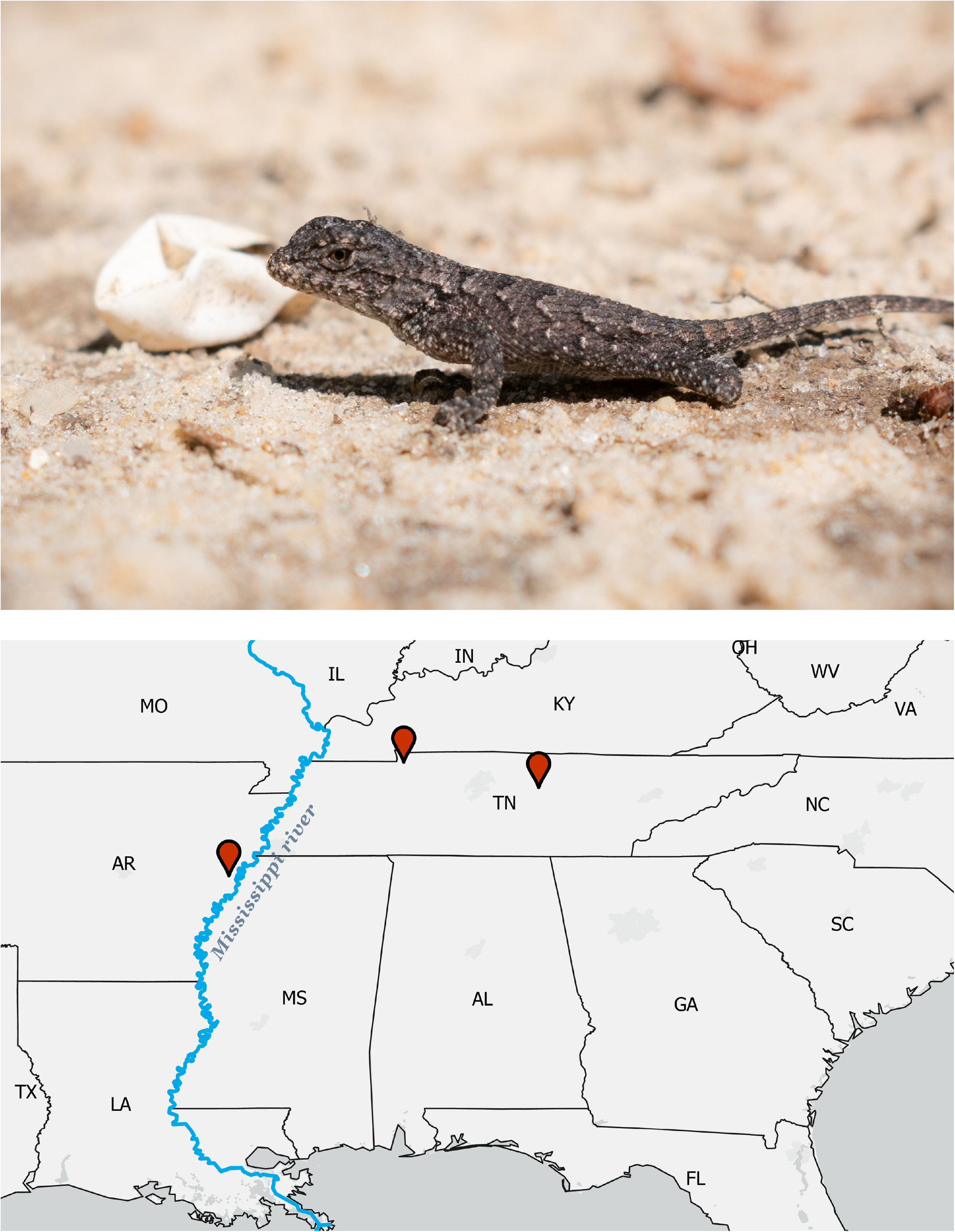
Above: Eastern fence lizard, *Sceloporus undulatus*, after hatching (individual not part of this study). Below: Maternal collection sites in Tennessee (*East*) and Arkansas (*West*).

## Materials and Methods

### Animal and microbial sampling

Lizard eggs were obtained from mothers in natural populations in Arkansas (AR) and Tennessee (TN), USA (Figure 1; hereafter, *West* and *East*, respectively). Mothers were housed from 12 to 22 days before ovipositing, and eggs were incubated and hatched in soil from a common source in an animal housing facility. Five to six non-siblings from the same population were housed together in common garden conditions. At 21 days of age, lizards went through a gradual shift in diet from fruit flies (*Drosophila melanogaster*) to house crickets (*Acheta domesticus*) as they grew in size, and were fed house crickets exclusively at 60 days of age onwards for three times per week. Room temperature was maintained at 23°C with a light cycle of 12:12, in addition to a heat lamp maintained daily from 08:00 to 16:00 to allow thermoregulation. Further details on animal collection and husbandry are provided elsewhere (Assis et al. 2021). All procedures were reviewed and approved by The Pennsylvania State University’s Institutional Animal Care and Use Committee (IACUC).

We used sterile swabs (FLOQSwabs® 516CS01 Ultra Minitip Flocked Swabs, Copan Diagnostics Inc., Murrieta, CA) to collect microbial samples from the cloacae of twenty lizards (eight females and five males from *West*, five females and two males from *East*) born from ten mothers (seven from *West*, three from *East*). Samples of this size have been demonstrated to be sufficient for capturing interpopulation differences in teleost fishes (Panteli et al. 2020). Specimens had been preserved together in a glass container with 70% ethanol within 24 hours of natural death at varying ages (range: 53 to 280 days) throughout the common garden period between September 2016 and May 2017. As part of the rearing protocol, lizards were under frequent veterinary observation and no illness was diagnosed during the housing period. Although leveraging preserved specimens may bring some limitations (see *Discussion*), it can still provide relevant information on microbial compositions while reducing the toll on natural populations. Microbial sampling of preserved specimens via cloacal swabbing has been validated in birds and reptiles, and shown to yield microbial representation comparable to those from fecal samples (Bassis et al. 2017; Vogtmann et al. 2017; Berlow et al. 2020; Bodawatta et al. 2020; but see Videvall et al. 2018). A study with *Mus musculus* has demonstrated that delayed sampling after death did not affect gut microbiome composition (Čížková et al. 2021). Still, in this study we are unable to assess how the circumstances surrounding individual’s death may have affected its microbiome. Finally, all specimens were sexually immature, eliminating the effect that copulation may have on microbial transmission in lizards (White et al. 2011).

### DNA extraction and amplification

Swab tips were excised using sterile scissors and were subject to DNA extraction using the NucleoSpin Soil DNA extraction kit (Macherey-Nagel Inc., Allentown, PA; catalogue: 740780) as per the manufacturer’s instructions. A sterility control of an unused excised swab tip was included, as well as a DNA extraction negative process control (*i*.*e*. no input sample). Bacterial composition was characterized with amplicon sequencing of the 16S rRNA gene (515F and 806R). The PCR mixes for both reactions were as follows: 12 µL of Platinum II Hot-Start PCR Master Mix, 1.5 µL of each primer (10 µM), 5 µL template DNA and 10 µL molecular grade water for a final PCR volume of 30 µL. Bacterial 16S rRNA gene PCR cycling conditions were as follows: 3 minutes at 94°C, 25 cycles of: 45 seconds at 94°C, 60 seconds at 50°C and 90 seconds at 72°C, and a final elongation step of 10 minutes at 72°C. PCR negative controls were included. The sterility and process controls yielded no detectable DNA, and the sterility controls, process controls and the PCR negative controls had no observable amplicons.

### Amplicon library preparation

Generated amplicons were first cleaned using Mag-Bind TotalPure NGS magnetic beads (Omega Bio-Tek Inc., Norcross, GA; catalogue: M1378-01). Cleaned amplicons were indexed with the following PCR ingredients: 12.5 µL of Platinum II Hot-Start PCR Master Mix, 2.5 µL of each index (10 µM) and 2.5 µL of sterile water for a final volume of 25 µL. The indexing PCR cycling conditions were as follows: 1 minute at 98°C, 8 cycles of: 15 seconds at 98°C, 30 seconds at 55°C and 20 seconds at 72°C, and a final elongation step of 5 minutes at 72°C. Sample normalization of the indexed amplicons was performed with the SequalPrep normalization plate kit (ThermoFisher Scientific, Waltham, MA; catalogue: A1051001). Normalized samples were pooled, concentrated with a CentriVap micro IR Vacuum Concentrator (Labconco, Kansas City, MO), and gel purified with the PureLink quick gel extraction kit (ThermoFisher Scientific; catalogue: K210012). The pooled library was sequenced on the Illumina MiSeq sequencing platform (2 × 250bp). Raw data files in FASTQ format were deposited in the NCBI sequence read archive under BioProject ID PRJNA874042.

### Sequence analysis

Raw demultiplexed 16S rRNA gene data were processed using the Quantitative Insights into Microbial Ecology pipeline QIIME 2 version 2021.11 (Caporaso et al. 2010). Paired-ended 16S rRNA gene sequences were trimmed and denoised using DADA2 (Callahan et al. 2016). Taxonomy was assigned using the classify-sklearn qiime feature classifier against the Silva v138 database (Quast et al. 2013) at the single nucleotide threshold (Amplicon Sequence Variants, ASVs). Sequences assigned as chloroplasts or mitochondria were removed, and ASVs with less than 45 sequences (0.01 %; 16S rRNA) were also removed. The cleaned 16S rRNA gene data were then rarefied at 49,720 sequences per sample.

### Statistical analysis

We used linear models to analyze the effects of age and maternal population on bacterial richness (Chao1), evenness (Pielou), and diversity (Shannon’s). For each metric, we began with a fully specified model containing a two-way interaction between the predictors along with its main effects, and followed with the removal of the interaction term if non-significant. We inspected models for overdispersion and highly influential data points (Cook’s Distance), which we did not observe (all < 0.52). Multicollinearity between main effects was estimated using the Variance Inflation Factor with the vif function in the R package *car* (Fox and Weisberg 2019). For all models in this study, all VIF < 4.18.

We evaluated bacterial compositional differences using a distance-based Redundancy Analysis (db-RDA) based on Bray-Curtis dissimilarity, as well as weighted UniFrac distances, using Permutational Multivariate Analyses of Variance (PERMANOVAs) with the adonis function in the R package *vegan*, along with their respective analyses of dispersion. Independent variables were age at sampling and maternal population along with an interaction between the two. The interaction was non-significant, so only the two main effects were retained in further analyses.

To identify changes in the relative abundance of specific taxa according to age, we first used data summarized at the genus level in a Similarity Percentages test (simper) from the package *vegan* based on Bray-Curtis dissimilarities of binary age classes. We used this test as a guide for individual genera to be investigated independently in downstream analyses, and for this reason did not adjust the significance level for multiple testing. Genera with p < 0.05 were analyzed separately using a Negative Binomial Generalized Linear Model (NBGLM) with host age as the predictor (α = 0.01).

## Results

The interaction between age and lineage approached a significant relationship with richness of ASVs: Adj R^2^ = 0.33, β = 0.35, p = 0.053. Negative coefficients for the main effects indicate a trend towards reduced bacterial richness with increasing age, with distinct slopes for each lineage. Removal from the model of the marginally non-significant interaction term resulted in a marginally nonsignificant relationship for individual age, with older individuals trending towards reduced bacterial richness (Adj R^2^ = 0.20, β = −0.19, p = 0.052). Bacterial evenness exhibited a significant association with an Age-by-Population interaction (Adj R^2^ = 0.30, β = −0.001, p = 0.04), in which the Western lineage showed decreasing evenness with individual age, a relationship not seen for the Eastern lineage. Finally, bacterial diversity had a significant relationship with individual age (Adj R^2^ = 0.35, β = −0.005, p < 0.01; Figure 2) after removal of the non-significant interaction term with population: for the two lineages, older individuals had significantly reduced indices of bacterial diversity. Even though the two youngest lizards in the sample (53 and 54 days) still had fruit flies as a minor component of their diet at the time of preservation before transitioning to exclusively crickets at 60 days, these two data points were not overly influential in driving these relationships (Figure 2; all Cook’s Distance < 0.52).

**Figure 2:**
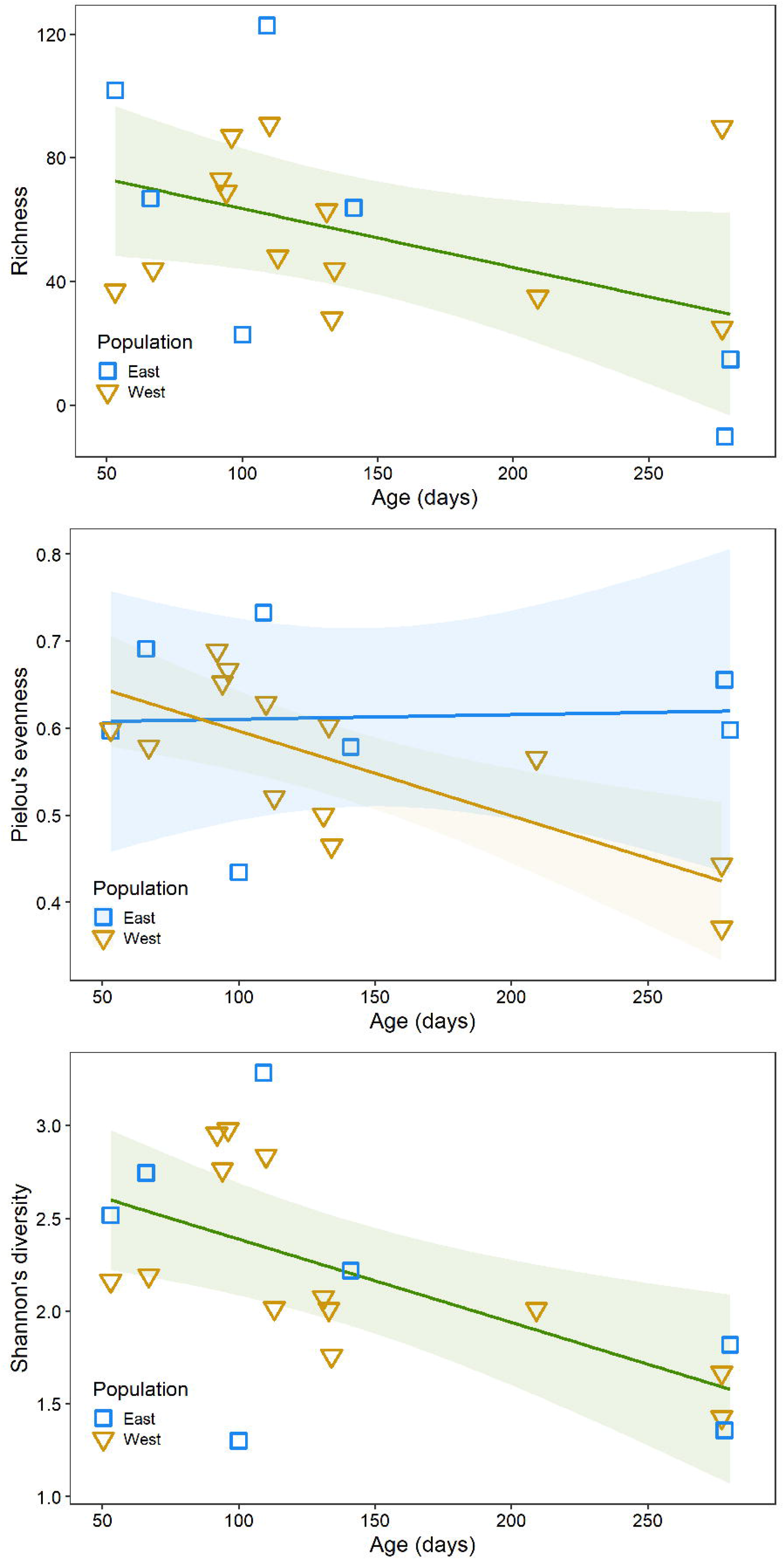
Linear relationships between lizard age at sampling and bacterial community richness (marginally nonsignificant, Adj. R^2^ = 0.20, β = −0.19, df = 17, p = 0.052), evenness (population-by-age interaction: Adj. R^2^ = 0.30, β < −0.01, df = 16, p = 0.045), and diversity (Adj. R^2^ = 0.35, β < −0.01, df = 17, p < 0.01). Gray bands represent 95% confidence intervals.

While contrasting bacterial composition between groups, we identified a significant effect of individual age when corrected for phylogenetic relationships (Figure 3a; PERMANOVA on weighted UniFrac: R^2^ = 0.11, F_1,19_ = 2.40, p = 0.003) and on db-RDA (Figure 3b; PERMANOVA: R^2^ = 0.12, F_1,19_ = 2.49, p = 0.004). No effect was seen for maternal population on microbiome composition (all p > 0.2), and analyses of dispersion did not suggest heterogeneous dispersion (all p > 0.4). Visual inspection of UniFrac and db-RDA plots indicates that with increasing age, bacterial communities diverged among individuals, even within lineages (Figure 3a, 3b).

**Figure 3:**
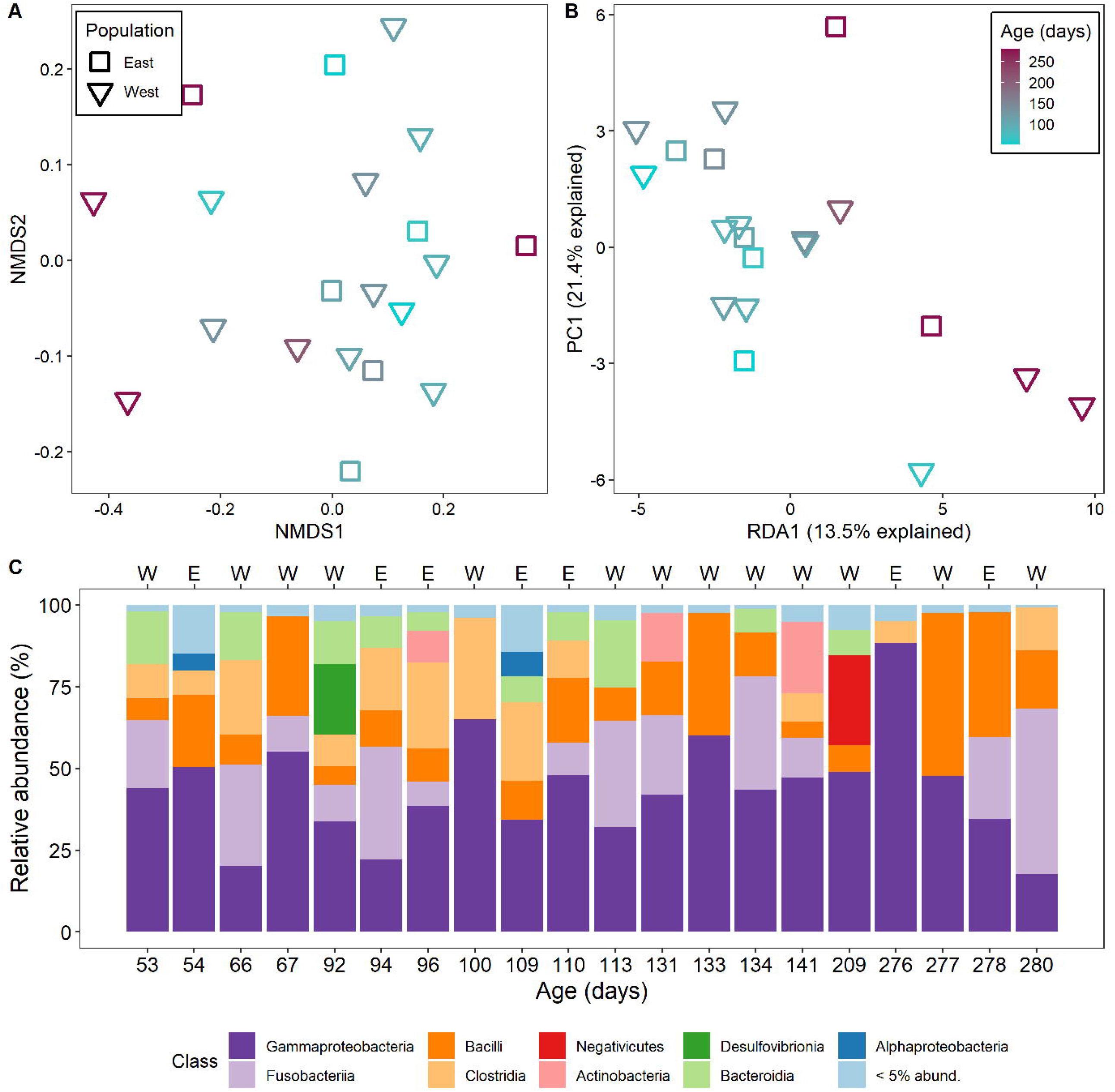
**A**: Weighted UniFrac distances of cloacal bacterial composition; stress level = 0.14. **B**: Distance-based Redundancy Analysis plot constrained on individual age and based on Bray-Curtis dissimilarities. **C**: Bacterial taxonomy plots with relative abundances summarized at the Class level. Communities are ordered according to lizard age at sampling. Letters above indicate maternal population (E: East; W; West).

As individual age had a significant influence on bacterial composition, we examined bacterial genera with significant shifts in abundance with individual age. Similarity Percentages analysis indicated 14 bacterial genera with contrasting abundances between age classes (p < 0.05). Seven of those showed acceptable NBGLM diagnostics (Cook’s distance < 1) and a significantly negative relationship with age (all z < −2.6; all p < 0.01). Those genera were: *Proteus, Clostridium_sensu_stricto_1, Paraclostridium, Vagococcus, Dysgonomonas*, an uncharacterized genus in the *Vermiphilaceae* family, and *Breznakia* (table S1). Between lineages, nine genera had contrasting abundances, but only two showed significance in NBGLM: the Western lineage had reduced abundances of *Mycobacterium* (z = −2.046 p = 0.04) and *Lactobacillus* (z = −3.563, p < 0.001).

## Discussion

*Sceloporus undulatus* lizards hatched from mothers collected East and West of the Mississippi river and reared in a laboratory common garden had indistinguishable cloacal bacterial composition and diversity, despite long withstanding allopatry between lineages (Leaché 2009; Wiens et al. 2010). Previous studies have identified strong environmental effects on microbiome composition between allopatric conspecifics (Moeller et al. 2017; Liu et al. 2021), including within the *Sceloporus* genus (Montoya-Ciriaco et al. 2020). However, we did not observe such differences when allopatric lineages of *S. undulatus* were reared under standardized laboratory conditions. Importantly, lizards sampled at older ages had distinct bacterial composition, indicating that longer exposure to the environment has a stronger impact on the microbiome than host genetics or maternal transmission, a pattern previously seen in the humans (Shaw et al. 2017; Rothschild et al. 2018). Notably, bacterial diversity steadily decreased in older individuals, akin to evenness for western lizards. Therefore, a homogeneous habitat and diet had a continuous, depreciative effect on diversity that was shared by both lineages but affected by evenness in only one of them. Analogous effects of captivity on microbiome diversity have been seen in other vertebrates such as horses (Metcalf et al. 2017), Nile tilapia (Deng et al. 2021), and the brown Kiwi (San Juan et al. 2021; but see McKenzie et al. 2017). Opposite treatments and effects have also been demonstrated: captive animals reintroduced into natural populations or fed natural diets can exhibit increased microbiome diversity (Chong et al. 2019; Martínez-Mota et al. 2020). We discuss the significance of these patterns in detail below.

Bacterial composition shifted with host age, but these shifts did not converge onto similar community structures. Instead, they became more distinct and less diverse among older individuals, even if from the same lineage. It is unclear whether the patterns seen here could also occur in wild individuals exposed to heterogeneous environments, or what the role of vertical transmission from maternal microbiomes could be. Maternal bacterial transmission appears to occur through inoculation of yolk prior to egg shelling, and these may be critical protectants against fungal infections in the embryo (Trevelline et al. 2018; Bunker, Elliott, et al. 2021; but see Van Veelen et al. 2018). In *S. undulatus*, bacterial composition in gravid females is distinct from non-gravid females (Trevelline et al. 2019) and maternal exposure to stress may cause analogous effects (MacLeod et al. 2022). Nevertheless, the lack of effect for maternal population seen here indicates reduced significance of vertical transmission in face of environmental exposure.

With increasing age, we detected a significant decrease in the relative abundance of bacteria assigned to the genera *Proteus, Clostridium_sensu_stricto_1, Paraclostridium, Vagococcus, Dysgonomonas*, an uncharacterized genus in the Vermiphilaceae family and *Breznakia*. A significant decrease in the relative abundance of these genera could be due to the interaction of different factors, including microbiome maturation and changing inter-microbial interactions with age (Stephens et al. 2016; Iozzo and Sanguinetti 2018). In a different study with captive crocodile lizards, bacteria assigned to the *Paraclostridium* genus had a reduced relative abundance relative to wild lizards, although no change with age was seen (Tang et al. 2020). ASVs assigned to the *Breznakia* and *Dysgonomonas* genera have previously been associated with the digestive systems of insects (Tegtmeier et al. 2016; Bridges and Gage 2021). Therefore, their presence could be due to the insect-based diet of the lizards and are likely transient rather than actual gut colonizers. A decrease in their relative abundance could indicate differences in digestion efficacy with age, but this requires further investigation.

Given the importance that gut microbiomes have in physiological and behavioral processes of the host (Vuong et al. 2017; Lynch and Hsiao 2019), these findings reinforce the importance of considering microbiome effects in laboratory experiments with captive animals and in the extrapolation of patterns to the natural environment, as has been prompted by others (Kohl et al. 2014; Ericsson and Franklin 2021; Zhang et al. 2022). Still, limitations in our experiment may affect the generalization of this conclusion. Because individuals were preserved, we were unable to quantify within-individual temporal changes in microbial composition, and thus rely on an assumption of equivalent baseline composition at hatching. Some support for this assumption may be found in the lack of population effects (this study) and in the uniformity of housing conditions and diet of all animals. It is also possible that individuals in nature experience analogous changes in microbiome composition as they age, and this should be a subject of future investigation. Still, the pattern seen here is well aligned with what has been seen in other systems and treatments in natural and controlled environments (Metcalf et al. 2017; Chong et al. 2019; Martínez-Mota et al. 2020; Deng et al. 2021; San Juan et al. 2021) A separate point is that cloacal swabs may provide only a portion of the whole microbiome picture, as different regions of the gut may have distinct compositions that could be driven differently (Bunker, Martin, et al. 2021). Still, some evidence suggests that qualitative differences may be conserved across gut regions or sampling methods in lizards (Kohl et al. 2017).

Our study indicates that a homogeneous environment and diet were linked to a continuous decrease of diversity in the bacterial microbiome of eastern fence lizards. No broad effects of lineage were observed, and thus no evidence for maternal population effects in shaping the cloacal microbiome of fence lizards differently. Overall, these findings improve our understanding of the genetic and environmental factors underlying the gut microbiome of oviparous vertebrates, while highlighting important subjects of future investigation.

## Notes

### Competing Interest Statement

The authors have declared no competing interest.

## References

Assis BA, Avery JD, Tylan C, Engler HI, Earley RL, Langkilde T. 2021. Honest signals and sexual conflict: Female lizards carry undesirable indicators of quality. Ecol Evol. 11:7647–7659. doi:10.1002/ece3.7598.

Bassis CM, Moore NM, Lolans K, Seekatz AM, Weinstein RA, Young VB, Hayden MK. 2017. Comparison of stool versus rectal swab samples and storage conditions on bacterial community profiles. BMC Microbiol. 17. doi:10.1186/s12866-017-0983-9.

Berlow M, Kohl KD, Derryberry EP. 2020. Evaluation of non-lethal gut microbiome sampling methods in a passerine bird. Ibis (Lond 1859). 162:911–923. doi:10.1111/ibi.12807.

Bodawatta KH, Puzejova K, Sam K, Poulsen M, Jønsson KA. 2020. Cloacal swabs and alcohol bird specimens are good proxies for compositional analyses of gut microbial communities of Great tits (Parus major). Anim Microbiome. 2. doi:10.1186/s42523-020-00026-8.

Bonder MJ, Kurilshikov A, Tigchelaar EF, Mujagic Z, Imhann F, Vila AV, Deelen P, Vatanen T, Schirmer M, Smeekens SP, et al. 2016. The effect of host genetics on the gut microbiome. Nat Genet. 48:1407–1412. doi:10.1038/ng.3663.

Bridges CM, Gage DJ. 2021. Development and application of aerobic, chemically defined media for Dysgonomonas. Anaerobe. 67. doi:10.1016/j.anaerobe.2020.102302.

Bunker ME, Elliott G, Heyer-Gray H, Martin MO, Arnold AE, Weiss SL. 2021. Vertically transmitted microbiome protects eggs from fungal infection and egg failure. Anim Microbiome. 3. doi:10.1186/s42523-021-00104-5.

Bunker ME, Martin MO, Weiss SL. 2021. Recovered microbiome of an oviparous lizard differs across gut and reproductive tissues, cloacal swabs, and faeces. Mol Ecol Resour. doi:10.1111/1755-0998.13573.

Callahan BJ, McMurdie PJ, Rosen MJ, Han AW, Johnson AJA, Holmes SP. 2016. DADA2: High-resolution sample inference from Illumina amplicon data. Nat Methods. 13:581–583. doi:10.1038/nmeth.3869.

Caporaso JG, Kuczynski J, Stombaugh J, Bittinger K, Bushman FD, Costello EK, Fierer N, Peña AG, Goodrich JK, Gordon JI, et al. 2010. QIIME allows analysis of high-throughput community sequencing data. Nat Methods. 7:335–336. doi:10.1038/nmeth.f.303.

Chong R, Grueber CE, Fox S, Wise P, Barrs VR, Hogg CJ, Belov K. 2019. Looking like the locals - gut microbiome changes post-release in an endangered species. Anim Microbiome. 1. doi:10.1186/s42523-019-0012-4.

Čížková D, Ľudovít Ď, Piálek J, Kreisinger J. 2021. Experimental validation of small mammal gut microbiota sampling from faeces and from the caecum after death. Heredity (Edinb). 127:141–150.

Clavel T, Gomes-Neto JC, Lagkouvardos I, Ramer-Tait AE. 2017. Deciphering interactions between the gut microbiota and the immune system via microbial cultivation and minimal microbiomes. Immunol Rev. 279:8–22. doi:10.1111/imr.12578.

Deng Y, Kokou F, Eding EH, Verdegem MCJ. 2021. Impact of early-life rearing history on gut microbiome succession and performance of Nile tilapia. Anim Microbiome. 3. doi:10.1186/s42523-021-00145-w.

Ericsson AC, Franklin CL. 2021. The gut microbiome of laboratory mice: considerations and best practices for translational research. Mamm Genome. 32:239–250. doi:10.1007/s00335-021-09863-7.

Fox J, Weisberg S. 2019. An {R} Companion to Applied Regression, Second Edition. Sage.

Gould AW, Zhang V, Lamberti L, Jones EW, Obadia B, Korasidis N, Gavryushkin A, Carlson JM, Beerenwinkel N, Ludington WB. 2018. Microbiome interactions shape host fitness. Proc Natl Acad Sci U S A. 115:E11951–E11960. doi:10.5061/dryad.2sr6316.

Iozzo P, Sanguinetti E. 2018. Early dietary patterns and microbiota development: still a way to go from descriptive interactions to health-relevant solutions. Front Nutr. 5. doi:10.3389/fnut.2018.00005.

Kohl KD, Brun A, Magallanes M, Brinkerhoff J, Laspiur A, Acosta JC, Caviedes-Vidal E, Bordenstein SR. 2017. Gut microbial ecology of lizards: insights into diversity in the wild, effects of captivity, variation across gut regions and transmission. Mol Ecol. 26:1175–1189. doi:10.1111/mec.13921.

Kohl KD, Skopec MM, Dearing MD. 2014. Captivity results in disparate loss of gut microbial diversity in closely related hosts. Conserv Physiol. 2. doi:10.1093/conphys/cou009.

Kreznar JH, Keller MP, Traeger LL, Rabaglia ME, Schueler KL, Stapleton DS, Zhao W, Vivas EI, Yandell BS, Broman AT, et al. 2017. Host genotype and gut microbiome modulate insulin secretion and diet-induced metabolic phenotypes. Cell Rep. 18:1739–1750. doi:10.1016/j.celrep.2017.01.062.

Leaché AD. 2009. Species tree discordance traces to phylogeographic clade boundaries in North American fence lizards (Sceloporus). Syst Biol. 58:547–559. doi:10.1093/sysbio/syp057.

Ley RE, Lozupone CA, Hamady M, Knight R, Gordon JI. 2008. Worlds within worlds: evolution of the vertebrate gut microbiota. Nat Rev Microbiol. 6:776–788. doi:https://doi.org/10.1038/nrmicro1978.

Liu R, Shi J, Shultz S, Guo D, Liu D. 2021. Fecal bacterial community of allopatric Przewalski’s gazelles and their sympatric relatives. Front Microbiol. 12. doi:10.3389/fmicb.2021.737042.

Lynch JB, Hsiao EY. 2019. Microbiomes as sources of emergent host phenotypes. Science. 365:1405–1409. doi:https://www.science.org/doi/10.1126/science.aay0240.

Macke E, Tasiemski A, Massol F, Callens M, Decaestecker E. 2017. Life history and eco-evolutionary dynamics in light of the gut microbiota. Oikos. 126:508–531. doi:10.1111/oik.03900.

MacLeod KJ, Kohl KD, Trevelline BK, Langkilde T. 2022. Context-dependent effects of glucocorticoids on the lizard gutmicrobiome. Mol Ecol. 31:185–196. doi:https://doi.org/10.1111/mec.16229.

Martínez-Mota R, Kohl KD, Orr TJ, Denise Dearing M. 2020. Natural diets promote retention of the native gut microbiota in captive rodents. ISME J. 14:67–78. doi:10.1038/s41396-019-0497-6.

McKenzie VJ, Song SJ, Delsuc F, Prest TL, Oliverio AM, Korpita TM, Alexiev A, Amato KR, Metcalf JL, Kowalewski M, et al. 2017. The effects of captivity on the mammalian gut microbiome. In: Integrative and Comparative Biology. Vol. 57. Oxford University Press. p. 690–704.

Metcalf JL, Song SJ, Morton JT, Weiss S, Seguin-Orlando A, Joly F, Feh C, Taberlet P, Coissac E, Amir A, et al. 2017. Evaluating the impact of domestication and captivity on the horse gut microbiome. Sci Rep. 7. doi:10.1038/s41598-017-15375-9.

Moeller AH, Suzuki TA, Lin D, Lacey EA, Wasser SK, Nachman MW. 2017. Dispersal limitation promotes the diversification of the mammalian gut microbiota. Proc Natl Acad Sci U S A. 114:13768–13773. doi:10.1073/pnas.1700122114.

Montoya-Ciriaco N, Gómez-Acata S, Muñoz-Arenas LC, Dendooven L, Estrada-Torres A, Díaz De La Vega-Pérez AH, Navarro-Noya YE. 2020. Dietary effects on gut microbiota of the mesquite lizard Sceloporus grammicus (Wiegmann, 1828) across different altitudes. Microbiome. 8. doi:10.1186/s40168-020-0783-6.

Newsome SD, Feeser KL, Bradley CJ, Wolf C, Takacs-Vesbach C, Fogel ML. 2020. Isotopic and genetic methods reveal the role of the gut microbiome in mammalian host essential amino acid metabolism. Proc R Soc B Biol Sci. 287:21192995. doi:10.1098/rspb.2019.2995.

Panteli N, Mastoraki M, Nikouli E, Lazarina M, Antonopoulou E, Kormas KA. 2020. Imprinting statistically sound conclusions for gut microbiota in comparative animal studies: A case study with diet and teleost fishes. Comp Biochem Physiol - Part D Genomics Proteomics. 36. doi:10.1016/j.cbd.2020.100738.

Parata L, Mazumder D, Sammut J, Egan S. 2020. Diet type influences the gut microbiome and nutrient assimilation of Genetically Improved Farmed Tilapia (Oreochromis niloticus). PLoS One. 15:e0237775. doi:10.1371/journal.pone.0237775.

Parata L, Nielsen S, Xing X, Thomas T, Egan S, Vergés A. 2020. Age, gut location and diet impact the gut microbiome of a tropical herbivorous surgeonfish. FEMS Microbiol Ecol. 96. doi:10.1093/femsec/fiz179.

Quast C, Pruesse E, Yilmaz P, Gerken J, Schweer T, Yarza P, Peplies J, Glöckner FO. 2013. The SILVA ribosomal RNA gene database project: improved data processing and web-based tools. Nucleic Acids Res. 41:D590–D596. doi:10.1093/nar/gks1219.

Rothschild D, Weissbrod O, Barkan E, Kurilshikov A, Korem T, Zeevi D, Costea PI, Godneva A, Kalka IN, Bar N, et al. 2018. Environment dominates over host genetics in shaping human gut microbiota. Nature. 555:210–215. doi:10.1038/nature25973.

Round JL, Mazmanian SK. 2009. The gut microbiota shapes intestinal immune responses during health and disease. Nat Rev Immunol. 9:313–323. doi:10.1038/nri2515.

San Juan PA, Castro I, Dhami MK. 2021. Captivity reduces diversity and shifts composition of the Brown Kiwi microbiome. Anim Microbiome. 3. doi:10.1186/s42523-021-00109-0.

Shaw L, Ribeiro ALR, Levine AP, Pontikos N, Balloux F, Segal AW, Roberts AP, Smith AM. 2017. The human salivary microbiome is shaped by shared environment rather than genetics: evidence from a large family of closely related individuals. MBio. 8:e01237–17. doi:https://doi.org/10.1128/mBio.01237-17.

Spor A, Koren O, Ley R. 2011. Unravelling the effects of the environment and host genotype on the gut microbiome. Nat Rev Microbiol. 9:279–290. doi:10.1038/nrmicro2540.

Stephens WZ, Burns AR, Stagaman K, Wong S, Rawls JF, Guillemin K, Bohannan BJM. 2016. The composition of the zebrafish intestinal microbial community varies across development. ISME J. 10:644–654. doi:10.1038/ismej.2015.140.

Sylvia KE, Demas GE. 2018. A gut feeling: microbiome-brain-immune interactions modulate social and affective behaviors. Horm Behav. 99:41–49. doi:10.1016/j.yhbeh.2018.02.001.

Tang GS, Liang XX, Yang MY, Wang TT, Chen JP, D. WG, Li H, Sun BJ. 2020. Captivity influences gut microbiota in crocodile lizards (Shinisaurus crocodilurus). Front Microbiol. 11. doi:10.3389/fmicb.2020.00550.

Tegtmeier D, Riese C, Geissinger O, Radek R, Brune A. 2016. Breznakia blatticola gen. nov. sp. nov. and Breznakia pachnodae sp. nov., two fermenting bacteria isolated from insect guts, and emended description of the family Erysipelotrichaceae. Syst Appl Microbiol. 39:319–329. doi:10.1016/j.syapm.2016.05.003.

Trevelline BK, MacLeod KJ, Knutie SA, Langkilde T, Kohl KD. 2018. In ovo microbial communities: A potential mechanism for the initial acquisition of gut microbiota among oviparous birds and lizards. Biol Lett. 14. doi:10.1098/rsbl.2018.0225.

Trevelline BK, Macleod KJ, Langkilde T, Kohl KD. 2019. Gestation alters the gut microbiota of an oviparous lizard. FEMS Microbiol Ecol. 95. doi:10.1093/femsec/fiz086.

Van Veelen HPJ, Salles JF, Tieleman BI. 2018. Microbiome assembly of avian eggshells and their potential as transgenerational carriers of maternal microbiota. ISME J. 12:1375–1388. doi:10.1038/s41396-018-0067-3.

Videvall E, Strandh M, Engelbrecht A, Cloete S, Cornwallis CK. 2018. Measuring the gut microbiome in birds: Comparison of faecal and cloacal sampling. Mol Ecol Resour. 18:424–434. doi:10.1111/1755-0998.12744.

Vogtmann E, Chen J, Amir A, Shi J, Abnet CC, Nelson H, Knight R, Chia N, Sinha R. 2017. Comparison of Collection Methods for Fecal Samples in Microbiome Studies. In: American Journal of Epidemiology. Vol. 185. Oxford University Press. p. 115–123.

Vuong HE, Yano JM, Fung TC, Hsiao EY. 2017. The microbiome and host behavior. Annu Rev Neurosci. 40:21–49. doi:10.1146/annurev-neuro-072116.

White J, Richard M, Massot M, Meylan S. 2011. Cloacal bacterial diversity increases with multiple mates: Evidence of sexual transmission in female common lizards. PLoS One. 6. doi:10.1371/journal.pone.0022339.

Wiens JJ, Kuczynski CA, Arif S, Reeder TW. 2010. Phylogenetic relationships of phrynosomatid lizards based on nuclear and mitochondrial data, and a revised phylogeny for Sceloporus. Mol Phylogenet Evol. 54:150–161. doi:10.1016/j.ympev.2009.09.008.

Wu Y, Yang Y, Cao L, Yin H, Xu M, Wang Z, Liu Y, Wang X, Deng Y. 2018. Habitat environments impacted the gut microbiome of long-distance migratory swan geese but central species conserved. Sci Rep. 8. doi:10.1038/s41598-018-31731-9.

Youngblut ND, Reischer GH, Walters W, Schuster N, Walzer C, Stalder G, Ley RE, Farnleitner AH. 2019. Host diet and evolutionary history explain different aspects of gut microbiome diversity among vertebrate clades. Nat Commun. 10. doi:10.1038/s41467-019-10191-3.

Zhang L, Yang F, Li T, Dayananda B, Lin L, Lin C. 2022. Lessons from the diet: Captivity and sex shape the gut microbiota in an oviparous lizard (Calotes versicolor). Ecol Evol. 12. doi:10.1002/ece3.8586.

